# Meteorological conditions influence short-term survival and dispersal in a reinforced long-lived bird population.

**DOI:** 10.1101/003228

**Authors:** Loïc A. Hardouin, A. Robert, M. Nevoux, O. Gimenex, F. Lacroix, Yves Hingrat

## Abstract

1. A high immediate mortality rate of released animals is an important cause of translocation failure (“release cost”). Post-release dispersal (i.e. the movements from the release site to the first breeding site) has recently been identified as another source of local translocation failure. In spite of their potential effects on conservation program outcomes, little is known about the quantitative effects of these two sources of translocation failure and their interactions with environmental factors and management designs.
2. Based on long-term monitoring data of captive-bred North African houbara bustards (hereafter, houbaras) over large spatial scales, we investigated the relative effects of release (e.g., release group size, period of release), individual (e.g., sex and body condition) and meteorological (e.g., temperature and rainfall) conditions on post-release survival (N = 957 houbaras) and dispersal (N = 436 houbaras).
3. We found that (i) rainfall and ambient air temperature had respectively a negative and a positive effect on houbara post-release dispersal distance, (ii) in interaction with the release period, harsh meteorological conditions had negative impact on the survival of houbaras, (iii) density dependent processes influenced the pattern of departure from the release site and (iv) post-release dispersal distance was male-biased, as natal dispersal of wild birds (although the dispersal patterns and movements may be influenced by different processes in captive-bred and in wild birds).
4. *Synthesis and applications.* Overall, our results demonstrate that post-release dispersal and mortality costs in translocated species may be mediated by meteorological factors, which in turn can be buffered by the release method. As the consequences of translocation programs on population dynamics depend primarily upon release costs and colonisation process, we suggest that their potential interactions with meteorological conditions be carefully addressed in future programs.

## Introduction

Although high immediate mortality of released animals is an important cause of translocation failure (“release cost”, Tavecchia et al. 2009), post-release dispersal (an analogue of natal dispersal, defined here as the movements from the release site to the first breeding site) was identified as another source of local failure (Le Gouar et al. 2008). At the local (population) scale, translocation may serve to enhance population growth or improve population viability (Seddon et al. 2012). At larger scales, such as the metapopulation scale, translocation may promote the connectivity of local populations and/or extend the species' range (Seddon et al. 2012). Although dispersal is a key component of metapopulation dynamics, it is also associated with local failure (Mihoub et al. 2011). Furthermore, if dispersal entails survival costs (Bonte et al. 2012), it may partially explain patterns of mortality associated with restoration projects (Tweed et al. 2003). Overall, the factors affecting mortality and dispersal following release events strongly influence the success or failure of population management and conservation strategies (Le Gouar et al. 2008, Mihoub et al. 2011, Oro et al. 2011).

While adequate post-release monitoring are now prevalent (Seddon et al. 2007), temporal and spatial replicates of release events are still uncommon (Le Gouar et al. 2008). Consequently, conservationists rely primarily on short-term survival and dispersal data to assess the effects of release on population dynamics and to inform release strategies. However, the short-term survival of a released population might poorly reflect persistence over the long-term (Armstrong et al. 1999). Moreover, individual decisions and abilities along the dispersal process are differently affected by proximate factors (Clobert et al. 2009). In particular, if variations in dispersal patterns and movements are shaped by the individual phenotype and environmental conditions, this may result in different dispersal patterns and movements in captive-bred and wild individuals, having in turn an impact on translocation success (Clobert et al. 2009). Hereafter, we will refer to the three stages of dispersal following Bonte et al. (2012)’s general definitions and nomenclature for wild populations (i.e., departure from natal area, transfer –the movement *per se* and settlement at the reproduction site), although adapted to the case of translocations (i.e., considering departure from the release site, see Le Gouar et al. 2012). Assessment of the multiple factors affecting survival and dispersal over different temporal and spatial scales is necessary for conservationists to comply with the objectives of the program (e.g. population persistence or connectivity among populations) and to determine the sustainability of translocation over time (Armstrong & Seddon 2008).

Post-release survival and dispersal of individuals may be influenced by two main types of factors: 1) those relating to the initial conditions of the release (for instance and non-exhaustively: release group size, period of release; hereafter, release factors) and 2) those relating to individual characteristics (hereafter, individual factors), such as sex, age, and body condition. Additionally, although it is likely that local meteorological factors can also greatly influence the survival and dispersal of translocated animals, they have, to our knowledge, never been investigated.

This paper evaluates the importance of short- and long-term processes and the relative effects of release, individual and local meteorological factors on the survival and dispersal of a captive-bred species in a reinforcement (i.e. an addition of individuals to an existing population, IUCN 2012) conservation program. We benefited from an important conservation and release effort involving long-term and large spatial-scale monitoring of captive-bred North African houbara bustards (*Chlamydotis undulata undulata*, Family Otididae; hereafter, houbaras) in Morocco.

In numerous species, (i) meteorological factors strongly influence individual survival (Nevoux et al. 2008), particularly in low quality, weak or inexperienced individuals (Robert et al. 2012); and (ii) released individuals suffer high immediate mortality after release (Tavecchia et al. 2009). Based on these two empirical findings, we predict that survival of released houbaras will be negatively affected by harsh meteorological factors (such as abundant rainfall and/or low temperature) and that their effects will be strongest shortly after release due to the inexperience of houbaras in their release habitat.

Although most theoretical predictions deal with dispersal rate but not with the timing of dispersal or the dispersal distance, most classical hypotheses on dispersal rates in patchy habitats were generalized to movement patterns and can be applied to non-colonial species and continuous habitat (e.g. Débarre & Gandon 2010). Accordingly, we predict that the time elapsed between release and departure will be negatively correlated with release group size, a surrogate of the local density at the time of release, as (i) the houbara’s release protocol involves very large groups and (ii) intraspecific competition has been identified as an important ultimate cause of dispersal in both theoretical works and empirical studies of many species (Perrin & Mazalov 2000), including houbara bustards (Hardouin et al. 2012). In addition, recent evidence suggests that natal dispersal distance is male-biased in this species (Hardouin et al. 2012). Therefore, if sex-specific dispersal behaviour is not altered by captive-breeding or release protocols and conditions, we predict that post-release dispersal distance of 115 houbaras will be male-biased.

## Methods

### GENERAL METHODS

#### Study sites and biological model

The North African houbara bustard is a non-migratory, sexually dimorphic species, with males being larger and heavier than females. It exhibits an exploded-lek mating system, where display males are less tightly aggregated than in a classical lek (Hingrat et al. 2008). This vulnerable species (BirdLife International 2012) has suffered severe declines in wild populations over the last 40 years. Conservation concern led in 1995 to the establishment of the Emirates Centre for Wildlife Propagation (ECWP) in Morocco (Lacroix 2003). A comprehensive strategy combining ecological research, reinforcement using captive-bred houbaras and hunting management was established in order to restore houbara populations and allow regulated traditional Arab falconry. A partnership was established between ECWP and Moroccan authorities for the conservation and management of houbara populations over an area of 74 400 km^2^ (Fig. 1). This study area represents between 8 and 13% of the species distribution range in North Africa (using Species Distribution Modelling with houbara presence thresholds of 0.4 and 0.5, A.-C. Monnet, unpublished data). Hunting was banned between 2000 and spring 2005 and then restricted to the wintering period (i.e. October to January) in delimited hunted areas totalizing 60% of the managed area. The present study was conducted from March 2001 to February 2010, then comprising a non-hunted period (2001 to spring 2005) and a hunted period (autumn 2005 to 2010).

**Figure 1:**
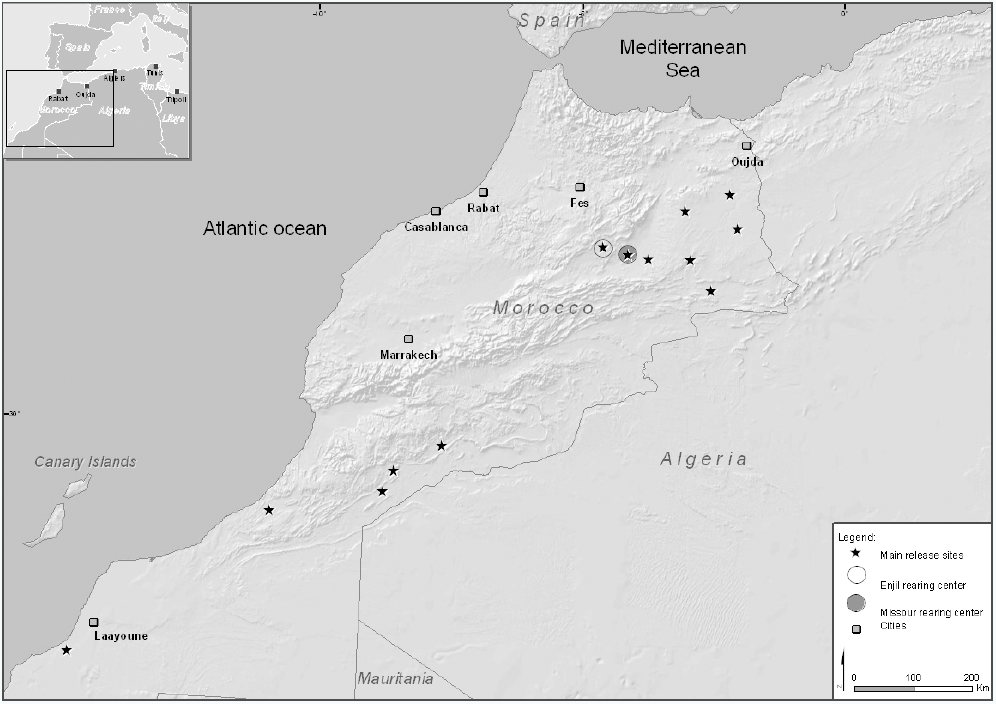
Locations of main release sites in Morocco. Houbaras were bred at two breeding stations: Missour (grey circle) and Enjil (white circle).

#### Breeding stations, release and radio-tracking procedure

Two ECWP breeding stations were established: one in Missour in 1995 and one in Enjil (Fig. 1) in 2005. Between 1998 and 2012, the ECWP has released 61,555 houbaras in North Africa, primarily in Morocco (see Lesobre et al. 2008 for insemination and rearing protocols, housing and pre-release conditions). Houbaras were released during two periods: from September to November (hereafter, “autumn release”) and from March to May (hereafter, “spring release”). In autumn, houbaras are mainly released in non-hunted areas to avoid direct mortality and disturbance due to hunting. The releases varied by sites (Fig. 1) and group size (from 10 ± 3 to 292 ± 101 houbaras per release) over time. Before release, each individual was weighed (± 1 g), blood-sampled (for molecular sexing) and tagged with a metal ring or subcutaneous electronic micro-chip (TROVAN LID100, DorsetID).

One thousand and four hundred houbaras (out of the 20,652 houbaras released during the study period) were fitted with necklace battery-powered transmitters with mortality signals (11g model RI-2B-M, 20g model RI-2D-M, Holohil System Ltd.). Sixty-five birds were recaptured before their batteries died and fitted with new 20g transmitters. Houbara bustards were monitored from the ground and by aerial telemetry (details in Hardouin et al. 2012). Birds were primarily ground-based monitored at their last location within a radius buffer of 5km. When not detected, birds were searched for using aerial telemetry (detection up to 80km, 303 ± 138 flying hours per year) for six months following an approximate radius of 40km from its last location and were systematically searched for during other aerial telemetry procedures (i.e. when searching for other “lost” birds). After six months without any location, a bird is considered as “lost” and searching is ceased. Locations were taken with GPS (accuracy ± 5 meters for terrestrial telemetry and ± 200 meters for aerial telemetry).

Two standard monitoring protocols were applied: 1) from their release to 90 days (± 10 days) post-release, houbaras were searched for twice a week over the entire study area in Morocco (Fig. 1), and 2) after 90 days, houbaras were searched for once a week. The first protocol was primarily used to get accurate measures of movement to analyse the departure and transfer phases of the short-term post-release dispersal (see below). Both protocols were used in combination to determine the distance from the release site to the first breeding site (i.e. settlement distance) and to build capture histories for survival analyses. For survival analysis, we considered only individuals released in north-eastern Morocco (Fig. 1), as release monitoring in the remaining study areas ceased after 90 days.

### POST-RELEASE DISPERSAL

#### Definition of movement parameters and transfer phase

Short-term post-release dispersal behaviour, defined as departure and transfer within three months of release of houbaras, was described with two distinct metrics.

Metric 1 corresponds to the linear distance between an individual’s release location and any of its successive locations; it measures the propensity to disperse away from the release site (Hardouin et al. 2012). Repeated measures of this metric over the whole monitoring period characterized the net dispersal distance (NDD). Metric 2 is the distance between two successive locations. From this metric, we calculated cumulative distance (CD), as the full path distances covered by an individual from the release site to the last location (Hardouin et al. 2012). This metric was used to estimate departure date (see below). Distances were measured using ArcGIS 9.2 (Environmental Systems Research Institute) and Hawth’s Analysis Tools (Beyer 2004). Only birds located at least 10 times and at least once per 30 days were considered for analysis. Birds that died or were lost during this period were excluded from analysis (Hardouin et al. 2012). Therefore, 436 individuals and 8,679 localizations were considered in this analysis, yielding 20 ± 8 (Mean ± SD) localizations, 92 ± 8 days of monitoring per individual and 4.7 ± 4.3 days between localizations.

#### Departure phase

The date of departure from the release site was estimated using cumulative distances (CD). Departure date was defined as the date at which the distance from the release sites begins to increase with time, inducing a break on the regression line between cumulative distances and time (detailed procedure in Appendix S1).

#### Settlement distance

Eighty six houbaras (out of the 436 individuals) were continuously monitored from their release until their first breeding attempt. Their settlement distance was estimated as the linear distance from the release site to the first nest (for females) or display site (for males). Released females initiated reproduction at 1.6 ± 0.5 years, whereas released males started displaying at 2.1 ± 0.8 years (data from our study sample), making them difficult to monitor over the longterm. We monitored 69 females and 17 males with respect to settlement distance.

#### Local meteorological data

In the north-eastern region of the study area, a network of 10 weather stations was installed close to the primary release sites (sFig. S1). Ambient air temperature (°C) and rainfall (mm) were recorded daily. For each bird’s location, we calculated the mean ambient air temperature and sum rainfall for the ± 3 days surrounding the localization event, using data from the nearest weather station. Rainfall and temperature were negatively correlated (r = −0.13, P < 0.001) with high temperature associated with dryness. We assessed the effects of local meteorological factors on houbaras during transfer using only individuals for which release site and subsequent localisations were near the weather stations to properly assess the effects of the meteorological variables (N = 223 out of 436 individuals). The mean distance between bird’s locations (N = 3,628 localizations) and weather stations was 17.8 ± 12.7 km; and did not exceed 65 km.

#### Released bird body condition and age

The 436 houbaras considered in the analysis were released at an average age of 234 ± 89 days, weighing 1,092 ± 140 g (females) and 1,482 ± 235 g (males) (see Table S1 for details). An index of body condition was extracted for each individual using the residuals of the regression of the body weight and release age (as body weight linearly increased with age at the time of the release; inflexion point of the growth curve = 44 ± 8 days for males and 38 ± 6 days for females) with sex as a covariate (F_2,433_ = 322.7, Adj.R^2^ = 0.6, P < 0.001, Hardouin et al. 2012).

#### Statistical analysis

We first investigated the effects of release and individual factors on departure date using linear mixed effect models. The model tested for effects of body condition (and its quadratic term), sex, release age, release group size and first order interactions on departure date. ‘Year’, ‘release site’ and ‘period’ were added as random terms. We then tested the effects of body condition (and its quadratic term), sex, release age, release group size, time after release (i.e., based on the dates of the localization events) and first order interactions on the NDDs to analyse the pattern of movement along the short-term post-release dispersal. We considered several random terms: ‘individual ID’ (nested into 'release site', N = 22), ‘year’ and ‘period of release’. Third, we tested for effects of local meteorological factors (i.e., rainfall and ambient air temperature), sex, body condition and their first order interactions on NDDs on individuals for which accurate meteorological data were available (N = 223 individuals, see above) to analyse their effects on the short-term dispersal distance between each individual’s localizations. The random slope terms were added as in the previous model and ‘time after release’ was considered as a random intercept to account for the repeated measures over time. Finally, we investigated whether settlement distance was explained by body condition at the time of the release, sex or age at first reproduction (as longer-distance dispersers might reproduce later). ‘Year of reproduction’ was added as a random slope term. For all models, we started from the global model (all explanatory variables and first order interactions) and compared its performance with submodels from which non-significant terms were deleted one at a time (see details in Appendix S2). NDDs and departure dates were log-transformed and the cumulative distances were square-root-transformed to meet normality and homoscedasticity assumptions. The freeware R 2.10.1 (R Development Core Team 2009) and the libraries lme4 (Bates & Maechler 2009), languageR (Baayen 2009) and segmented (Muggeo 2008) were used for statistical analyses.

### SURVIVAL ANALYSIS

We used multi-event capture-recapture modelling (Pradel 2005) using a mixture of alive recaptures and dead recoveries to estimate survival of radio-marked individuals (N = 957 individuals) from March 2001 to February 2010. Capture occasions were defined as a time-step of three months (N = 36 occasions). We studied the effect of sex and release period (autumn and spring) on short- (next occasion) and long-term (following occasions) survival probability according to years. The time dependent model was coded to possibly obtain (when relevant): 1) the survival in the third months after release (i.e. short-term survival) for each release cohort, sex and year and 2) the annual survival on the long–term (after the third months post-release) for each release cohort and sex. We also assessed the influence of individual and temporal covariates on survivorship. Each bird was monitored for an average of 351 ± 400 days (max = 2,721 days). Details of the sample used in this analysis and of the model structure are given in Table S2 & Appendix S3.

Briefly, we considered two states to code whether an individual is alive or dead and 3 events to code for the observed fate of an individual at each occasion (0: not observed, 1: alive individual, 2: recovery of a dead individual). We verified the fit of the general, time-dependent model with program U-CARE version 2.3.2 (Choquet et al. 2009a, details in Appendix S3). A step-down model selection was performed using program E-SURGE version 1.7.1 (Choquet et al 2009b) based on the Akaike Information Criterion corrected for sample size and overdispersion (QAICc). When ΔQAICc was smaller than 2, we selected the model with the smallest number of parameters, following the parsimony principle (Lebreton et al. 1992). Starting from the time-dependent model (Appendix S4), we thus proceeded of three successive steps of model selection: 1) we performed model selection for recovery probabilities (Models E1 to E10 in Table S3); 2) recapture probabilities (Models C1 to C4 in Table S4); 3) and survival probabilities (Models S1 to S16 in Table S5). Unless indicated otherwise, estimates of survival, detection and recovery probabilities are presented as the value ± standard error (S.E.).

#### Selection of individual and meteorological covariates and modelling

To test whether variation in meteorological conditions explains a significant part of the temporal variation in survival rate, we integrated meteorological information as temporal covariates into models. At the spatial scale of our study (Figs. 1 & S1), no data on temporal variation in the abundance of houbara food resources were available. However, the availability of plant and animal species in the houbara’s diet is known to vary considerably over space and time (Bourass et al. 2012), and meteorological parameters have been used to characterize the environmental fluctuations faced by individuals (Nevoux et al. 2008). We used the daily temperature and rainfall data from the 10 weather stations (Fig. S1) to create annual meteorological indices. We then calculated the annual mean daily temperature (ADT) and rainfall amount (ADR) over the ten stations. After 6 years of reinforcement, regulated hunting on houbaras occurred from 2005 to 2009 in the winter (non-breeding) season. Although hunting was not our focus and autumn releases mainly occurred in non-hunted areas, we accounted for hunting in our analyses by integrating a temporal covariate describing the presence (1) or absence (0) of hunting in a given year in our model with meteorological covariates.

We first assessed the effects of meteorological indices on the selected time-dependent model to estimate which one (ADT, ADR or their interaction) had the greatest effect on temporal variation in survival (Models T1 to T11 in Table S6). The contribution of temporal covariates to the model was assessed using an analysis of deviance (ANODEV), and the amount of temporal variation in the focal rate explained by the covariate was evaluated by *R*^2^*dev* (Skalski et al. 1993).

We also tested for the effects of individual covariates describing the initial conditions of birds and group size upon release. We used an index of body condition (BC) (computed as above), the age of individuals at release (Ra) and the release group size (RGr) to assess whether these covariates influence survival rates. We individually tested each survival parameter of the time-dependent selected model, resulting in 16 candidate models with individual covariates (Models 1 to 16 in Table S7). A covariate is important when the associated 95% confidence interval (CI) of the corresponding slope does not include zero.

### RESULTS

#### DISPERSAL

##### Effects of release and individual factors on departure date

Departure dates were calculated for 396 of the 436 individuals included in this analysis and occurred on average 29.7 ± 20.2 days after release (median = 24.7, range = 2 - 90 days). From the selected model (R^2^ = 0.13, χ^2^_null_ = 8.2, df =3, P = 0.04), departure date was weakly but significantly explained by a positive interaction between body condition and release group size (MCMCmean= 1.04, HPD95 {1.001 − 1.08}, P = 0.04): individuals in poor body condition released in a large group tended to leave the release site first.

##### Effects of release and individual factors during transfer

The most parsimonious model of NDDs (R^2^ = 0.65, χ^2^_null_ = 2387.8, df = 7, P < 0.001) revealed significant effects of body condition, release age and time after release and first order interactions of time after release with body condition, release age and sex (Table 1). Individuals in better body condition moved farther distances than those in poorer body condition (Fig. 2). Older released birds travelled farther distances shortly after release, but not later (Table 1, Fig. S2). Moreover, the pattern of movement between males and females differed over time (sex * time after release interaction, Table 1). Males moved shorter distances than females shortly after release, but over time, the pattern reversed. This interaction suggests an increase of NDD in males or a decrease of NDD in females. Release group size did not affect NDDs (Release group: MCMCmean= 0.0007, HPD95 {−0.0006 − 0.002}, P = 0.4).

**Table 1:** Effects of release and individual factors on net dispersal distance in houbaras. The results of the selected linear mixed-effects model are given, with NDD as the response variable and body condition, sex, release age, time after release and their interactions as explanatory variables. Parameter estimates (i.e., fixed-effects estimates), HPD intervals, MCMCmeans and p-values based on the posterior distribution (pMCMC) for the fixed-effects table are given. For random effects, the estimates of the residual standard deviation, the standard deviation of the random effects (intercept), HPD intervals and MCMCmeans are provided. ‘Time’ refers to ‘time after release’.

| Fixed terms | Estimate | MCMCmean | HPD95lower | HPD95upper | pMCMC |
| --- | --- | --- | --- | --- | --- |
| Intercept | 1980.24 | 1923.25 | 391.00 | 10627.22 | <.001 |
| Body condition | 1.17 | 1.17 | 1.13 | 1.22 | <.001 |
| Sex | 0.93 | 0.92 | 0.82 | 1.04 | 0.19 |
| Release age | 1.18 | 1.20 | 1.06 | 1.35 | 0.002 |
| Time | 5.83 | 5.84 | 4.85 | 6.94 | <.001 |
| Body condition x Time | 0.9994 | 0.9994 | 0.9991 | 0.9998 | 0.001 |
| Sex x Time | 1.1339 | 1.1385 | 1.0167 | 1.2798 | 0.008 |
| Release age x Time | 0.9986 | 0.9986 | 0.9979 | 0.9992 | <.001 |

| Random terms | Std.Dev. | MCMCmean | HPD95lower | HPD95upper |
| --- | --- | --- | --- | --- |
| Individual : Release site | 0.69 | 0.49 | 0.46 | 0.52 |
| Release site | 0.36 | 0.33 | 0.23 | 0.45 |
| Year | 0.26 | 0.29 | 0.15 | 0.44 |
| Period of release | 0.15 | 0.76 | 0.01 | 1.57 |
| Residual | 0.75 | 0.77 | 0.76 | 0.78 |

**Figure 2:**
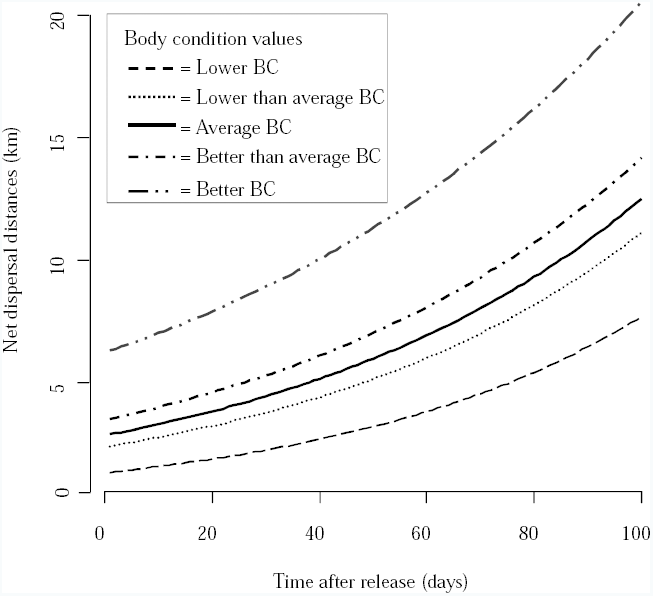
Partial plot representing the interaction between time and body condition (BC, given as quantiles) according to the net dispersal distance (in km) of houbaras (log-transformed data were back-transformed).

##### Effects of meteorological factors during transfer

According to the selected model (R^2^ = 0.76, χ^2^_null_ = 16.1, df = 3, P = 0.001), body condition was still positively associated with movements when including meteorological factors as explanatory variables (estimate ± SE = 0.0012 ± 0.0004, t-value = 3.1). Along monitoring, individuals covered greater distance when facing high ambient air temperature (estimate ± SE = 0.0133 ± 0.0045, t-value = 2.9) and low rainfall conditions (estimate ± SE = −0.0226 ± 0.0067, t-value = −3.3).

##### Effects of release and individual factors on settlement distances

The selected model (R^2^ = 0.21, df = 1, χ^2^_null_ = 7.15, P = 0.007) revealed that settlement distance was sex-biased, with males settling farther from the release site than females (MCMCmean = 0.61, HPD95 {0.15; 1.05}, P = 0.008). Neither body condition nor age at first reproduction significantly affected settlement distance.

#### SURVIVAL

Model selection procedures are detailed in Appendix S4, Tables S3 & S4 for encounter probabilities and Table S5 for survival.

##### Modelling variation in survival rates

Ultimately, the model [*S*_*a*1\**period*1\**t*+*a*1\**period*2+*a*2\**t*_] assuming short- vs. long-term- (i.e., a1 and a2, respectively), time- and release period-dependent survival was selected (Model S1 in Table S5). In particular, short-term survival differed according to the period of release (i.e., autumn vs. spring, period 1 vs. period 2, respectively), with time-dependence on short-term survival for individuals released in autumn ([*S*_*a*1\**period*1\**t*_] range between 0.19 ± 0.07 in 2009 and 0.92 ± 0.06 in 2006, Fig. 3a), while short-term survival was constant over years for releases occurring in spring ([*S*_*a*1\**period*2_] = 0.86 ± 0.02). Long-term survival varied according to year ([*S*_*a*2\**t*_], Fig. 3b), with a relatively low survival rate in 2008 (0.72 ± 0.03).

**Figure 3:**
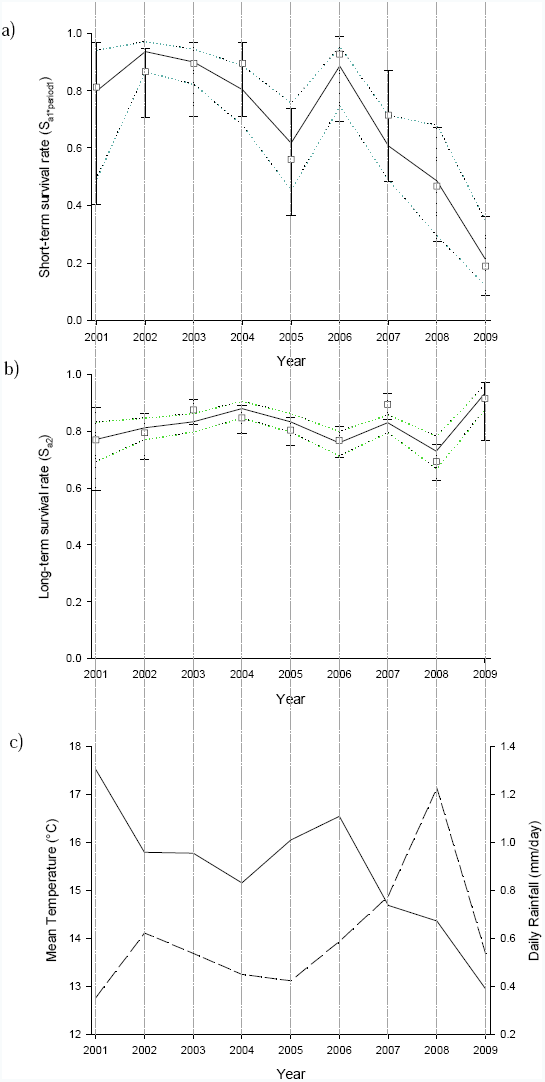
Annual variation in houbara survival rates: a) Survival rates estimated for the first time-class (i.e., the third month) and released in autumn, b) Survival rates for the second time-class (i.e., more than 3 months). For both, we present survival rates estimated from the time-dependent model and the 95% confidence intervals (white square). Survival rates (black lines) and 95% confident intervals (dotted lines) are estimated from the temporal covariates for a) with the interaction between ADT and ADR. [*S*_*a*1\**period*1\**t*\*(*ADT*\**ADR*)_] and for b) survival rates and 95% confident interval estimated with additive effect of ADT and ADR [*s*_*a*2\**t*\*(*ADT*+*ADR*)_]. c) Annual variation of the ambient air temperature (ADT, straight black line) and the daily rainfall (ADR, straight dashed line). Note that hunting stopped from winter 2001 to 2004 and started from winter 2005. The presence (1) / absence (0) of hunting were integrated in all the models as temporal covariate. Note that [S_*a*1\**period*2_] is constant = 0.86 ± 0.02 and not shown.

##### Effect of the meteorological covariates on survivorship

Meteorological covariates were incorporated into the selected time-dependent model [*S*_*a*1\**period*1\**t*+*a*1\**period*2+*a*2\**t*_] (Table S6). The interaction between annual temperature (ADT) and rainfall (ADR) explained a significant part of the short-term survival for autumn releases ([*S*_*a*1\**period*1\**t*\*(*ADT*\**ADR*+*hunting*)_], Fig. 3a), and an additive effect of ADT and ADR explained a significant part of the long-term survival ([*S*_*a*2\**t*\*(*ADT*+*ADR*+*hunting*)_], Fig. 3b, Models T1 in Table S6). The selected meteorological indices explained 83% of the deviance in the temporal variation of survival probabilities (F-Anodev_1,9_ = 44.6, P < 0.001). It is important to note that when hunting is removed from the previous model, 76% of the deviance was still explained by meteorological covariates (F-Anodev_1,11_ = 35.7, P < 0.001).

##### Effect of the individual covariates on survivorship

Individual covariates were incorporated into the selected time-dependent model (Table S7). The models with ‘Release age’ [*cov*_*Ra*_] as covariates (Models 1, 2 and 3 in Table S7) on long-term survival were preferred over all of the other models, with age at release being negatively related to the long-term survival rates ([*S*_*a*1\**period*1\**t*+*a*1\**period*2+*a*2\**t*+*a*2*(*i*+*cov*_*Ra*_)_]: slope = −0.20 ± 0.07; CI {−0.05; −0.35}, Model 1 in Table S7).

#### DISCUSSION

We found that (1) density-dependent processes (e.g. intra-specific competition) partly shaped the pattern of (early) post-release dispersal, as suggested by the interaction between initial density (release group size) and body condition; (2) post-release dispersal was male-biased, as in natal dispersal of wild houbaras (Hardouin et al. 2012); nevertheless, our results suggest that the dispersal decision may be influenced by different processes in wild and captive-bred birds; and (3) meteorological factors had critical effects on houbara survival, although these effects can be mitigated by appropriate timing/method of release.

##### Effects of the meteorological, release and individual factors on post-release dispersal phases

Our results highlight the importance of meteorological conditions during the transfer phase of post-release dispersal and their potential impact on the colonisation processes of translocated populations. We found that rainfall and ambient air temperature had a negative and a positive effect, respectively, on net dispersal distance, independently of the release period. These meteorological factors can have indirect effects mediated through food availability. Recent findings showed that the houbara plant diet is dominated by young green leaves of annual plants, whereas ants dominate the animal diet of the species (Bourass et al. 2012). The availability of these food resources is driven by ambient air temperature and rainfall. For instance, while rainfall induces instant new sprout and high ant foraging behaviour, warm conditions are associated with low plant availability and low ant activity (e.g. Bernstein 1975, Whitford 1978, López et al. 1993). Meteorological conditions can also directly affect flight ability, having in turn an impact on dispersal distances (Walls et al. 2005).

The release group size interacted with body condition and affected departure from the release site. This finding suggests that individuals with poor body condition leave the vicinity of the release site sooner in larger release groups. Birds in good condition may be less affected by density-dependent processes (such as intraspecific competition) before their departure than birds in poor body condition (Matthyssen 2005). These birds may remain at the release site for longer to limit the release cost and may gain sufficient reserve and/or experience in their novel environment. This behaviour may increase their chance to survive, as opposed to the birds in poor body condition that seem to leave the release site prematurely in this high-density context. Previous results on wild juvenile houbaras indicate that males in poor body condition move farther than those in good condition during natal dispersal; distance being independent of body condition in wild females (Hardouin et al. 2012). Here, captive-bred houbaras show the opposite pattern suggesting that selective pressures exerted by captivity (Robert 2009) and/or the release protocol (Le Gouar et al. 2012) may interact with the ultimate factors shaping individual dispersal strategies (e.g., inbreeding avoidance and intra-specific competition). Finally, in contrast to wild juveniles that already occupy favourable breeding habitats, released individuals may have to search more extensively for lekking or nest sites. Thus, only individuals in good condition may afford the distance-dependent cost of dispersal (Bonte et al. 2012). Hence, older released birds had longer net dispersal distances than younger ones just after release, but not later. Presumably, the longer distances initially covered by older individuals reflect differences in muscular development or hormonal activity such as an increase in corticosterone with age during captivity (Dufty & Belthoff 2001).

Finally, we detected male-biased dispersal in houbaras, consistent with studies of natal dispersal in wild houbaras (Hardouin et al. 2012). Sex-biased dispersal is generally interpreted in the theoretical frame of the inbreeding avoidance hypothesis (Szulkin et al. 2013). Recent findings in Morocco demonstrated a lack of kin association within and between leks in male houbara but a higher genetic relatedness in males than in females (Lesobre 2008). This finding suggests greater philopatry in males (Lesobre 2008), in contrast to our results. However, rare long-distance dispersal events in females (3 females dispersed more than 100 km in our study) suggest that they may be vectors of gene flow at the metapopulation level (Ibrahim et al. 1996). This hypothesis is also supported by our data on breeding dispersal distance (i.e. the distance between two successive breeding sites): males, once settled, remain near their display sites (mean breeding dispersal distance ± SD = 2.3 ± 4.6 km; median = 438 m; N = 16 males, unpublished data), whereas females exhibited farther breeding dispersal distances (mean = 16.4 ± 30.4 km, median = 2.9 km; N = 39 females, unpublished data).

##### Effects of the meteorological, release and individual factors on short- and long-term survival

Meteorological conditions strongly influence individual survival (e.g., Grosbois et al. 2008, Nevoux et al. 2008) and, consequently, population dynamics and viability (e.g., little bustards Tetrax tetrax, Morales et al. 2005). While several studies have reported substantial short-term mortality costs in translocated populations (Tavecchia et al. 2009), these costs have primarily been attributed to release methods, individual characteristics and habitat requirements. Here, we report a strong interaction between release strategy (autumn vs. spring releases) and meteorological conditions on short-term survival, which suggests that release costs are likely to be modulated by environmental stochasticity (Nicoll et al. 2003). Indeed, short-term survival was strongly associated with annual rainfall, ambient air temperature and their interaction, but only for individuals released in autumn. In particular, low temperatures in association with either flooding or aridity negatively affected short-term survival. In contrast, the short-term survival of individuals released in spring remained high and constant over years. Finally, we did not observe any difference in long-term survival between release cohorts. Annual rainfall and temperature also affected long-term survival, particularly in 2008, when severe flooding occurred in the study area. Although environmental stochasticity is in essence unpredictable, knowledge of its magnitude and its complex effects on the life histories of released individuals should greatly improve the accuracy of demographic projection models designed to examine the feasibility of translocation protocols. We thus advocate that environmental fluctuations be considered in future release strategies in houbaras and other translocation efforts (Robert et al. 2007, Sæther et al. 2007).

Our results also showed that younger released individuals are more tolerant later in life to meteorological variation. The choice of the release age in translocation program is particularly important and conservationists are often divided on favouring either age or habitat related advantages in captive-bred individuals (Sarrazin and Legendre 2000). In our context, all individuals might pay the cost of release, as no learning relevant to survival in the wild environment (e.g., identification of resource locations, anti-predator behaviour) can occur in pre-release aviaries. However, older released individuals might enjoy short-term advantages (e.g., in terms of competitive ability) and/or gain more experience with ageing that fade outside of the captive environment. Therefore, selection on older individuals might be expressed later, potentially explaining their higher mortality over the long-term. Time spent in captivity can alter the behaviour and/or physiology of individuals (e.g., Pedersen & Jeppesen 1990) in ways that negatively affect survival once released (Bremmer-Harrison et al. 2004). Further investigations are needed to understand the behavioural and/or physiological basis of age variation in survival. Finally, survival rates of captive-bred houbaras appear to approach or even exceed those of wild houbaras, although a survival analysis on the same time-scale with both wild and captive-bred houbaras would be necessary to fully evaluate this. Within three months of dispersal, wild juveniles (particularly females) exhibited overall lower survival rates (from 2006-2009: males = 0.86 ± 0.05 and females = 0.58 ± 0.06; Hardouin et al. 2012) than adults, and adult survival rates (from 2006-2009: males = 0.91 ± 0.02 and females = 0.83 ± 0.03) were similar those reported here.

##### Conservation implications and conclusions

In conservation translocations, post-release dispersal rate can be particularly high after release (Hester et al. 2008). Thus, understanding the factors influencing post-release dispersal is crucial in managing the trade-off between site fidelity and adaptive dispersal (Le Gouar et al. 2012). Whereas high post-release dispersal rates can be associated with dispersal related project failures (e.g. dispersal of released individuals outside the focal populations, Le Gouar et al. 2008), the absence of dispersal can limit habitat choice processes or be detrimental at larger (e.g. metapopulation) scale (Trakhtenbrot et al. 2005). Our results suggest that short-term post-release dispersal can be greatly influenced by captive breeding and release factors (i.e. dispersal patterns are distinct from those of wild-born individuals and post-release dispersal is influenced by timing and local release conditions) that however do not impact longer-term dispersal in terms of distance. Therefore, the assessment of the post-release dispersal at different temporal scales seems essential in translocated populations. While initial conditions lead to different responses between wild and captive-bred houbaras, these do not result in strong differences in natal and post-release dispersal distances (natal dispersal distance: wild male 19.4 ± 12.8 km, wild female = 8.6 ± 6.5 km; Hardouin et al. 2012). Moreover, sex-biased dispersal that constitutes an essential element in the connectivity and gene flow between populations occurs in both captive-bred and wild populations. Our results suggest that both the release group size and the period of release have to be carefully determined to limit premature departures of weaker individuals and long distance dispersal that may be induced by harsh meteorological conditions. Previous works have shown that local densities at the release site can also influence dispersal rate and pattern (Le Gouar et al. 2008). Further investigations on the effect of the local density could be helpful in refining release strategies. Overall, our results on the effects of the factors (including sex) influencing post-release dispersal distance enable general predictions of movement among individuals in translocated and natural populations. It is indeed important to account for post-release dispersal in population projections (Armstrong & Reynolds 2012) and to distinguish mortality from dispersal to adopt appropriate management strategies and maximise future establishment success (Tweed et al. 2003).

Our results also highlight the importance of long-term monitoring program to assess survival cost, as short-term survival rates do not accurately reflect long-term survival. Our findings further suggest a strong short-term sensitivity of individuals to unpredictable meteorological variation in autumn, in contrast to individuals released in spring, when meteorological conditions are generally milder. No subsequent effect of release period was found on long-term survival, suggesting that once individuals gathered experience in their novel environment, individuals from both release periods have similar survival probabilities. As the consequences of translocation programs on population dynamics depend primarily upon release costs, we suggest that their potential interactions with meteorological conditions be carefully addressed in future programs. Finally, and although survival and dispersal constitute critical components of population dynamics, recruitment and fertility rates need to be thoroughly assessed in future work to have a complete view of the demography of released houbaras. The Population Viability Analysis framework (Beissinger & McCullough 2002) represents the most useful approach to integrate the effects of meteorological fluctuations, hunting and management actions on the whole species life cycle. Such integration will provide reliable insights on the viability of the released population, and in turn, on the contribution of the conservation translocation program to the viability of the whole species.

## ACKNOWLEDGMENTS

Funding and supervision was provided by the Emirates Center for Wildlife Propagation (ECWP) under the leadership of the International Fund for Houbara Conservation (IFHC). We are grateful to H.H. Sheikh Mohammed bin Zayed Al Nahyan, Crown Prince of Abu Dhabi and Chairman of the IFHC and to H.E. Mohammed Al Bowardi, Deputy Chairman of the IFHC, for their support. Many thanks go to Jacques Renaud, General Manager of RENECO for Wildlife Preservation, for his support. We kindly thank all ECWP fieldworkers and breeders, Sylvain Boullenger, Eric le Nuz, Thibault Dieuleveut, Gwénaëlle Levêque, Chloé Deschamps, Guy Maurice, Vincent Lieron and Jean-François Léger for their contributions to data collection, their full commitment in the coordination of data collection, database and GIS management, and their helpful advice throughout this study. We are particularly indebted to Drs Anne-Christine Monnet, Marion Valeix, Simon Chamaillé-Jammes, Todd Katzner, Pierre Legagneux, Vincenzo Penteriani and Brian Preston for their helpful comments at different stages of the manuscript.

### DATA ACCESSIBILITY

The detailed survival model descriptions are available in the online supporting information and data files used in this paper would be available on http://datadryad.org/.

## Supporting Information

Additional supporting information may be found in the online version of this article

**Appendix S1.** Details of the breakpoint procedure.

**Table S1.** Details of the dispersal analysis sample of houbaras.

**Figure S1.** Map with the locations of the weather stations in the study area in Morocco.

**Appendix S2.** Model selection procedure of the linear mixed effect models.

**Table S2.** Details of the survival sample of houbaras.

**Appendix S3.** Goodness-of-fit and structure of the capture-recapture models.

**Figure S2.** Plot of the interaction between time and release age according to the NDDs of houbaras.

**Appendix S4.** Modelling variation in recovery and detection rates.

**Table S3.** Model selection on recovery probabilities.

**Table S4.** Model selection for detection rates.

**Table S5.** Model selection for survival rates.

**Table S6.** Model selection with temporal covariates.

**Table S7.** Model selection with individual covariates.

## REFERENCES

Armstrong, D.P., Castro, I., Alley, J.C., Feenstra, B. & Perott, J.K. (1999) Mortality and behaviour of hihi, an endangered New Zealand honeyeater, in the establishment phase following translocation. Biological conservation, 89, 329–339.

Armstrong, D.P. & Seddon, P.J. (2008) Directions in reintroduction biology. Trends in Ecology and Evolution, 23, 20–25.

Armstrong, D.P. & Reynolds, M.H. (2012). Modelling reintroduced population: the state of the art and future directions. Reintroduction Biology: integrating science and management (eds J.G. Ewen, D.P. Armstrong, K.A. Parker & P.J. Seddon), pp 165–222. Wiley-Blackwell, Oxford, UK.

Baayen, R.H. (2009) languageR: Data sets and functions with “Analyzing Linguistic Data: A practical introduction to statistics”. R package version 0.955.

Bates, D. & Maechler, M. (2010) lme4: linear mixed-eff ects models using S4 classes. R package ver. 0.999375-35.

Beissinger, S.R. & McCullough, D.R. (2002) Population Viability Analysis. University of Chicago Press, Chicago.

Bernstein, R.A. (1975) Foraging strategies of ants in response to variable food density. Ecology, 56, 213–219.

Beyer, H.L. (2004) Hawth’s Analysis Tools for ArcGIS. Available at http://www.spatialecology.com/htools.

BirdLife International (2012) Chlamydotis undulata. IUCN 2013. IUCN Red List of Threatened Species. Version 2013.1.

Bonte, D., Van Dyck, H., Bullock, J.M., Coulon, A., Delgado, M., Gibbs, M., Lehouck, V., Matthysen, E., Mustin, K., Saastamoinen, M., Schtickzelle, N., Stevens, V.M., Vandewoestijne, S., Baguette, M., Barton, K., Benton, T.G., Chaput-Bardy, A., Clobert, J., Dytham, C., Hovestadt, T., Meier, C.M., Palme, S.C.F., Turlure, C. & Travis, J.M.J. (2012) Costs of dispersal. Biological reviews, 87, 290–312.

Bourass, K., Léger, J.-F., Zaime, A., Qninba, A., Rguibi, H., El Agbani, M.A., Benhoussa, A. & Hingrat, Y. (2012) Observations on the diet of the North African houbara bustard during the non-breeding season. Journal of Arid Environment, 82, 53–59.

Bremmer-Harrison, S., Prodohl, P.A. & Elwood, R.W. (2004) Behavioural trait assessment as a release criterion: boldness predicts early death in a reintroduction programme of captive-bred swift fox (Vulpes velox). Animal Conservation, 7, 313–320.

Choquet, R., Lebreton, J.-D., Gimenez, O., Reboulet, A.-M. & Pradel, R. (2009a) U-CARE: Utilities for performing goodness of fit tests and manipulating Capture-Recapture data. Ecography, 32, 1071–1074.

Choquet, R., Rouan, L. & Pradel, R. (2009b) Program E-SURGE: a software application for fitting multievent models. Environmental and ecological statistics: modeling demographic processes in marked populations (eds D.L. Thomson, E.G. Cooch & M.J. Conroy), pp. 845–865. Springe-Verlag, New-York.

Clobert, J., Le Galliard, J.-F., Cote, J., Meylan, S. & Massot, M. (2009) Informed dispersal, heterogeneity in animal dispersal syndromes and the dynamics of spatially structured populations. Ecology Letters, 12, 197–209.

Débarre, F., & Gandon, S. (2010). Evolution of specialization in a spatially continuous environment. Journal of evolutionary biology, 23, 1090–1099.

Dufty, A.M.J. & Belthoff, J.R. (2001) Proximate mechanisms of natal dispersal: the role of body condition and hormones. Dispersal (eds J. Clobert, E. Danchin, A.A. Dhondt & J.D. Nichols), pp. 217–229. Oxford University Press: New York.

Grosbois, V, Gimenez, O Gaillard, J.-M., Pradel, R., Barbraud, C. Clobert, J. Møller, A.P. & Weimerskirch, H. (2008) Assessing the impact of climate variation on survival in vertebrate populations. Biological Reviews, 83, 357–99.

Hardouin, L.A., Nevoux, M., Robert, A., Gimenez, O., Lacroix, F. & Hingrat, Y. (2012) Determinants and costs of natal dispersal in a lekking species. Oikos, 121, 804–812.

Hester, J.M., Price, S.J., & Dorcas, M.E. (2008) Effects of relocation on movements and home ranges of eastern box turtles. The Journal of Wildlife Management, 72, 772–777.

Hingrat, Y., Saint Jalme, M., Chalah, T., Orhant, N. & Lacroix, F. (2008) Environmental and social constraints on breeding site selection. Does the exploded-lek and hotspot model apply to the Houbara bustard Chlamydotis undulata undulata? Journal of Avian Biology, 39, 393–404.

Ibrahim, K.M., Nichols, R.A. & Hewitt, G.M. (1996) Spatial patterns of genetic variations generated by different forms of dispersal during range expansion. Heredity, 77, 282–291.

IUCN (2012) Guidelines for reintroductions and other conservation translocations. IUCN, Gland, Switzerland. Available from www.issg.org/pdf/publications/Translocation-Guidelines-2012.pdf

Lacroix, F. (2003) The emirates center for wildlife propagation: developing a comprehensive strategy to secure a self-sustaining population of Houbara bustards in eastern Morocco. Houbara News, 5, 2.

Lebreton, J.D., Burnham, K.P., Clobert, J. & Anderson, D.R. (1992) Modeling survival and testing biological hypotheses using marked animals: a unified approach with case studies. Ecological monographs, 62, 67–118.

Le Gouar, P., Robert, A., Choisy, J.-P., Henriquet, S., Lecuyer, P., Tessier, C. & Sarrazin, F. (2008) Roles of survival and dispersal in reintroduction success of griffon vulture (gyps fulvus). Ecological Applications, 18, 859–872.

Le Gouar, P., Mihoub, J.-B. & Sarrazin, F. (2012) Dispersal and habitat selection: behavioural and spatial constraints for animal translocations. Reintroduction Biology: integrating science and management (eds J.G. Ewen, D.P. Armstrong, K.A. Parker & P.J. Seddon), pp. 138–164. Wiley-Blackwell, Oxford, UK.

Lesobre, L. (2008) Structures génétiques des populations menacées d’outardes houbaras (Chlamydotis undulata undulata). Implications à la gestion d’un élevage conservatoire et au renforcement des populations. PhD thesis, Muséum National d’Histoire Naturelle, Paris.

López, F., Acosta, F.J. & Serrano, J.M. (1993) Responses of the trunk routes of a harvester ant to plant density. Oecologia, 93, 109–113.

Matthyssen, E. (2005) Density-dependant dispersal in birds and mammals. Ecography, 28, 403–416.

Mihoub, J.-B., Robert, A., Le Gouar, P. & Sarrazin, F. (2011) Post-Release Dispersal in Animal Translocations: Social Attraction and the “Vacuum Effect”. PLoS ONE, 6(12), e27453.

Morales, M.B., Bretagnolle, V. & Arroyo, B. (2005) Viability of the endangered Little bustard Tetrax tetrax population of western France. Biodiversity and Conservation, 14, 3135–3150.

Muggeo, V.M.R. (2008) Segmented: an R package to fit regression models with broken-line relationships. Rnews, 8, 20–25.

Nevoux, M., Barbraud, J.-C. & Barbraud, C. (2008) Nonlinear impact of climate on survival in a migratory white stork population. Journal of Animal Ecology, 77, 1143–1152.

Nicoll, M.A.C., Jones, C., & Norris, K. (2003) Declining survival rates in a reintroduced population of the Mauritius kestrel: evidence for non-linear density dependence and environmental stochasticity. Journal of Animal Ecology, 72, 917–926.

Oro, D., Martínez Abraín, A., Villuendas, E., Sarzo, B., Mínguez, E., Carda, J., Alberdi, M. & Genovart, M. (2011) Lessons from a failed translocation program with a seabird species: determinants of success and conservation value. Biological Conservation, 144, 851–858.

Pedersen, V. & Jeppesen, L.L. (1990) Effects of early handling on later behaviour and stress responses in the silver fox (Vulpes vulpes). Applied Animal Behaviour Science, 26, 383–393.

Perrin, N. & Mazalov, V. (2000) Local competition, inbreeding, and the evolution of sexbiased dispersal. The American Naturalist, 155, 116–127.

Pradel, R. (2005) Multievent: an extension of multistate capture recapture models to uncertain states. Biometrics, 61, 442–447.

R Development Core Team (2009) R: A language and environment for statistical computing. R Foundation for Statistical Computing, Vienna, Austria. ISBN 3-900051-07-0, URL http://www.R-project.org.

Robert, A. (2009). Captive breeding genetics and reintroduction success. Biological Conservation, 142, 2915–2922.

Robert, A., Couvet, D. & Sarrazin, F. (2007) Integration of demography and genetics in population restorations. Ecoscience, 14, 463–471.

Robert, A., Paiva, V.H., Bolton, M., Jiguet, F. & Bried, J. (2012) The interaction between reproductive cost and individual quality is mediated by oceanic conditions in a long-lived bird. Ecology, 93, 1944–1952.

Sæther, B.E., Lillegård, M., Grøtan, V., Filli, F. & Engen, S. (2007) Predicting fluctuations of reintroduced ibex populations: the importance of density dependence, environmental stochasticity and uncertain population estimates. Journal of Animal Ecology, 76, 326–336.

Sarrazin, F., & Legendre, S. (2000). Demographic approach to releasing adults versus young in reintroductions. Conservation Biology, 14, 488–500.

Seddon, P.J., Armstrong, D. & Maloney, R.F. (2007) Developing the science of reintroduction biology. Conservation Biology, 21, 303–312.

Seddon, P.J., Strauss, W.M. & Innes, J. (2012) Animal translocations: what are they and why do we do them? Reintroduction Biology: integrating science and management (eds J.G. Ewen, D.P. Armstrong, K.A. Parker & P.J. Seddon), pp 1–32. Wiley-Blackwell, Oxford, UK.

Skalski, J.R., Hoffmann, A. & Smith, S.G. (1993) Testing the significance of individual and cohort-level covariates in animal survival studies. The use of marked individuals in the study of bird population dynamics: Models, methods, and software (eds J.D. Lebreton & P.M. North), pp 1–17. Birkhauser Verlag, Basel.

Szulkin, M., Stopher, K.V., Pemberton, J.M. & Reid, J.M. (2013) Inbreeding avoidance, tolerance, or preference in animals? Trends in ecology & evolution, 28, 205–211.

Tavecchia, G., Viedma, C., Martínez-Abraín, A., Bartolomé, M.-A., Gómez, J.A. & Oro, D. (2009) Maximizing re-introduction success: assessing the immediate cost of release in a threatened waterfowl. Biological conservation, 142, 3005–3012.

Trakhtenbrot, A., Nathan, R., Perry, G., & Richardson, D.M. (2005) The importance of long-distance dispersal in biodiversity conservation. Diversity and Distributions, 11, 173–181.

Tweed, E.J., Foster, J.T., Woodworth, B.L., Oesterle, P., Kuehler, C., Lieberman, A.A., Powers, A.T., Whitaker, K., Monahan, W.B., Kellerman, J. & Telfer, T. (2003) Survival, dispersal, and home-range establishment of reintroduced captive-bred puaiohi, Myadestes palmeri. Biological Conservation, 111, 1–9.

Walls, S.S., Kenward, R.E., & Holloway, G.J. (2005) Weather to disperse? Evidence that climatic conditions influence vertebrate dispersal. Journal of Animal Ecology, 74, 190–197.

Whitford, W.G. (1978) Foraging in seed-harvester ants Pogonomyrex spp. Ecology, 59, 185–189.

